# Developmental switching in *Physarum polycephalum*: Petri net analysis of single cell trajectories of gene expression indicates responsiveness and genetic plasticity of the Waddington quasipotential landscape

**DOI:** 10.1101/151878

**Authors:** Britta Werthmann, Wolfgang Marwan

## Abstract

The developmental switch to sporulation in *Physarum polycephalum* is a phytochrome-mediated far-red light-induced cell fate decision that synchronously encompasses the entire multinucleate plasmodial cell and is associated with extensive reprogramming of the transcriptome. By repeatedly taking samples of single cells after delivery of a light stimulus pulse, we analysed differential gene expression in two mutant strains and in a heterokaryon of the two strains all of which display a different propensity for making the cell fate decision. Multidimensional scaling of the gene expression data revealed individually different single cell trajectories eventually leading to sporulation. Characterization of the trajectories as walks through states of gene expression discretized by hierarchical clustering allowed the reconstruction of Petri nets that model and predict the observed behavior. Structural analyses of the Petri nets indicated stimulus- and genotype-dependence of both, single cell trajectories and of the quasipotential landscape through which these trajectories are taken. The Petri net-based approach to the analysis and decomposition of complex cellular responses and of complex mutant phenotypes may provide a scaffold for the data-driven reconstruction of causal molecular mechanisms that shape the topology of the quasipotential landscape.

## 1. Introduction

The organs and tissues of multicellular organisms are composed of different types of cells that are specialized in structure and function. These specialized cells are formed by a process called cell differentiation. The regulatory control of transcription factors leads to the differential expression of cell type-specific sets of genes encoding proteins that determine the morphology, the functional capabilities, and the behavior of the cells. Understanding the regulatory control of cell differentiation is of outstanding interest in basic research and especially in biomedicine, given the potential therapeutic applications of stem cells to tissue repair and the regeneration of organs or considering the dysregulation of proliferation and differentiation in cancer.

There are two basic and potentially complementing approaches in systems biology [1]. Bottom-up and top-down modeling. In the first case, models are built from known molecular interactions. In the second approach, also called reverse engineering or network inference, the model of the network is reconstructed based on experimental data. In principle, the quasipotential landscape controling the differentiation of individual cells could be calculated from a set of differential equations describing the biochemical reactions within the regulatory network [2, 3]. However, such a model is currently not available and information about the gene regulatory network is sketchy even for mammalian cells [3]. Computational analysis of known networks or network motifs has shown that their dynamic behaviour depends on mechanistic details, kinetic rate constants, and concentrations [4-7]. Given this situation and considering the complexity of cellular signaling, the reverse engineering (network inference) approach to complex processes like cell fate decisions may be a useful complement to bottom-up research. Hence, we wish to expore ways to reconstruct the quasipotential landscape from experimental time series data.

Cell differentiation is not just found in multicellular animals and plants. In many simple eukaryotes, specialized cell types occur in temporal order in the course of a life cycle, typically in response to environmental conditions that are sensed by specific receptor proteins [8]. *Physarum polyecephalum* is a classical model organisms in which cell differentiation processes have been studied early on.

For several reasons, the *P. polycephalum* plasmodium is a great object for studying differentiation at the single cell level [9]. The plasmodium is a giant single cell that can be grown to arbitrary size. The myriad of nuclei that are suspended in its cytoplasm display natural synchrony in cell cycle regulation and differentiation [10-15]. Because of this natural synchrony and because of the vigorous cytoplasmic shuttle streaming, the plasmodium provides a source from which macroscopic samples of homogeneous cell material can be repeatedly taken in order to analyse how the concentration of molecular components changes within a single cell as a function of time. To set a defined starting point, sporulation of a starving plasmodium can be triggered by a short pulse of far-red light which activates a phytochrome photoreceptor [16-18]. At around five to six hours after the pulse, the plasmodium becomes irreversibly committed to sporulation [9]. The formation of fruiting bodies starts at approximately eleven hours and the macroscopically visible morphogenesis is completed at about 18 hours after the pulse (Fig. 1). During the formation of the mature fruiting body, the protoplasm is cleaved and mononucleate, haploid spores are formed by meiosis.

**Figure 1.**
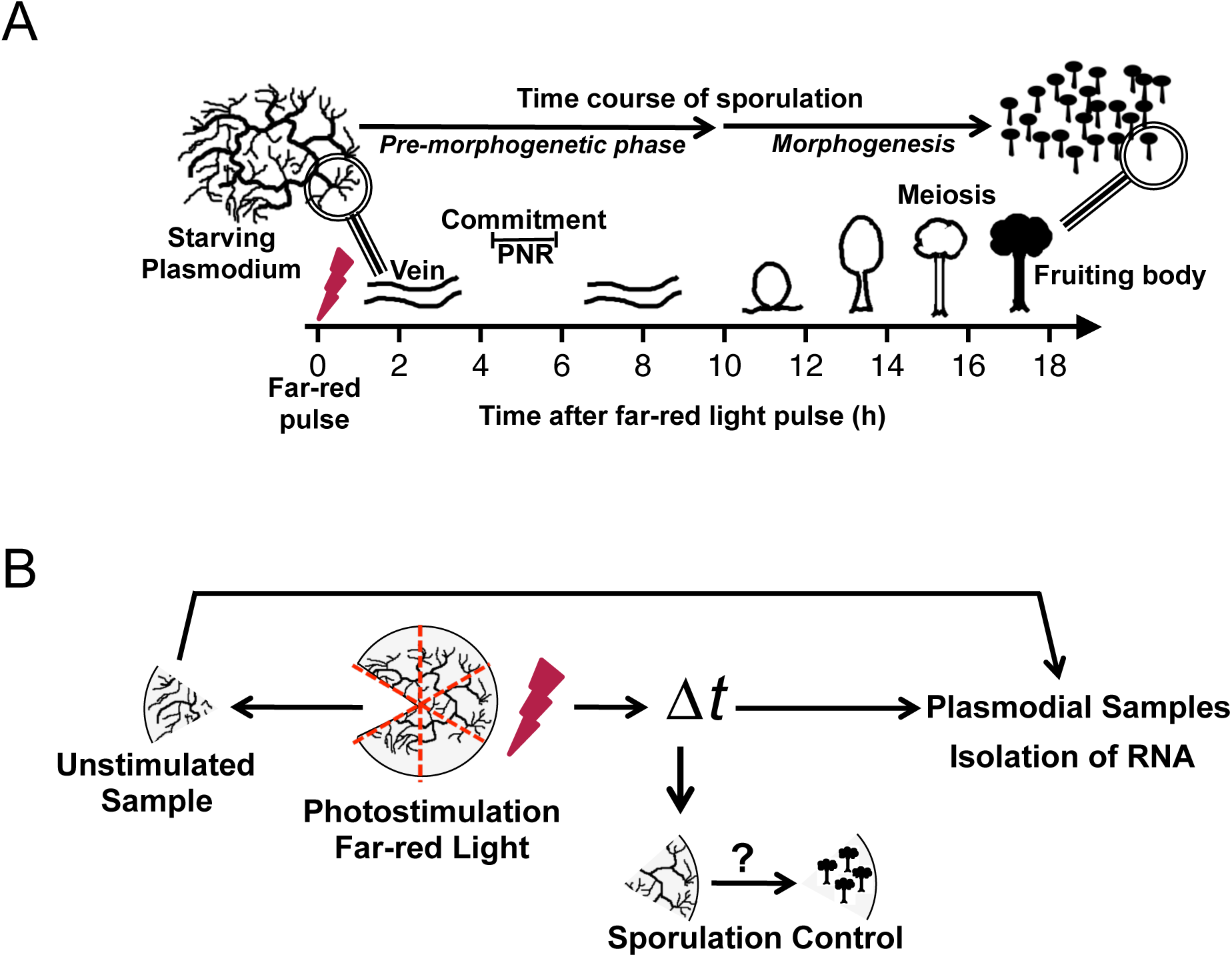
Time course of commitment and sporulation of a *P. polycephalum* plasmodium and preparation of samples for gene expression time series datasets. (A) Sporulation of a starving, competent, multinucleate (macro-) plasmodium can be triggered by a brief pulse of far-red light which is sensed by a phytochrome photoreceptor. After the inductive light pulse there is a pre-morphogenetic phase of about 9 to 10 hours without any visible changes in the plasmodial morphology. By crossing the point of no return (PNR) at four to six hours after induction, the plasmodium is committed to sporulation and loses its ablility to grow on nutrient agar. Morphogenesis starts when the plasmodial strands wind up and subsequently dissociate into small nodular structures (nodulation stage). Each nodule culminates and differentiates into a melanized fruiting body. Meiosis and cleavage of the multinucleate protoplasmic mass results in the formation of haploid, mononucleate spores that are released when the fruiting body ruptures. Under favourable conditions, a haploid, mononucleate amoeba can hatch from a spore to start a new life cycle [24]. (B) Collection of plasmodial samples for the measurement of gene expression time series. Starved plasmodial cells (each Petri dish contained one plasmodial cell on a plate of starvation agar) were stimulated with a pulse of far-red light and returned to the dark (for details see Materials and Methods). Before (referred to as 0h sample) and at various time points (2h, 6h, 8h, 11h) after the stimulus pulse, samples from the plasmodium were taken with a spatulum and shock frozen in liquid nitrogen for the subsequent isolation of RNA. The remainder of the plasmodium was further incubated until the next day to see whether the developmental decision to sporulation has occurred (sporulation control). For obtaining dark control time series, the experiment was performed exactly as described above, but the far-red light stimulus was omitted. Finally, the gene expression pattern in each sample was analysed.

Sporulation is associated with a cascade of differentially expressed genes [19-22]. By repeatedly taking samples at different time points after triggering sporulation with a far-red light pulse, we have shown that individual plasmodial cells proceed to sporulation along variable pathways that involve qualitatively different correlation patterns of differentially expressed genes [23]. Based on these intitial analyses we could however not specify routes that individual cells would take nor could we assign the differential regulation of specific genes to such routes.

Here, we identify trajectories of *P. polycephalum* plasmodial cells that had been exposed to a differentiation-inducing far-red light stimulus. Based on similarities and differences of single cell trajectories obtained by multidimensional scaling of gene expression data, we construct a phenotype model in the form of a Petri net that predicts the transition between states of gene expression discretized by hierarchical clustering. The Petri net formally links these states to the differential regulation of specific genes and to the sensory control of induction, commitment, and sporulation. Experimental comparison of mutants with different propensities to sporulate and structural analysis of the derived Petri nets indicates both, reponsiveness and genetic plasticity of the Waddington quasipotential landscape. We conclude that Petri nets that predict single cell trajectories by decomposing complex phenotypes may serve as formal scaffolds for the reconstruction (reverse engineering) of causal molecular mechanisms that define and shape the quasipotential landscape from experimental data.

## 2 Methods

### 2.1 Strains and growth of cell material

Cells of strains of PHO3 [25] and PHO57 [26] that were isolated in genetic screens for mutants with reduced propensity for light-induced sporulation were grown as microplasmodial suspensions in a fermenter for four days. Microplasmodia were harvested and washed, and the plasmodial pellet was allowed to dry on five layers of filter paper for 30 minutes. A ring of 1 g of the resulting cell paste was applied to the center of a 9 cm Ø Petri dish cantaining starvation agar using a motor-driven 50-mL syringe coupled to an in-house built automatic device for rotating the agar plate around its axis. Plasmodial cells were subsequently allowed to fuse on starvation agar so that a single macro-plasmodium developed on each plate. Macro-plasmodia were starved for 7 days in complete darkness at 22°C. Experimental conditions including media for growth and starvation were exactly as previously described [26]. Heterokaryons were obtained by applying 1 g of a 1:1 mixture of plasmodial pastes of strains PHO3 and PHO57 to the starvation agar plate and the cultures were further incubated as described above. During incubation, microplasmodia of the two strains spontaneously fused with each other to form a heterokoaryon.

### 2.2. Stimulation of plasmodial cells, preparation of samples, and gene expression analysis

Starved plasmodial cells (each Petri dish contained one plasmodial cell) were stimulated with a pulse of far-red light (*λ* ≥ 700 nm, 13 W/m^2^) as described (Starostzik & Marwan 1998) and returned to the dark (22°C). Before (referred to as 0 h sample) and at various time points (2 h, 6 h, 8 h, 11 h) after the stimulus pulse, samples from the plasmodium were taken with a spatula and shock frozen in liquid nitrogen for the subsequent isolation of RNA. The remainder of the plasmodium was further incubated until the next day and served as a sporulation control that indicated whether the developmental decision to sporulation occurred (Fig. 1B). Alternatively, time series were taken exactly as described while omitting the far-red light stimulus pulse (dark control time series). Total RNA was isolated from each sample separately, reverse transcribed, and the relative expression level of 35 marker genes (Supplementary Table 1) quantified by GeXP multiplex reverse transcription– polymerase chain reaction (RT–PCR) as previously described [26]. Primer sequences are described in [27]. One time series dataset was obtained for each plasmodial cell. A set of genes that showed differential regulation in response to far-red light stimulation in the wild type [27] were selected for further analysis (Table 1).

**Table 1.** Differentially regulated genes analysed for the reconstruction of single cell trajectories named according to their orthologs in the Uniprot database. Differential regulation is indicated for each gene as it occurs in cells committed to sporulation.

| Gene | Similarity | UniProt ID | E-value | Query coverage (%) | Differential regulation |
| --- | --- | --- | --- | --- | --- |
| *psgA* | *Physarum* specific gene | NA | NA | NA | down |
| *pldC* | Phospholipase D | Q9LRZ5 | 4E-14 | 61 | down |
| *pldB* | Phosphatidylinositol-glycan-specific phospholipase D | P80108 | 1E-80 | 83 | down |
| *pikB* | Phosphatidylinositol 3-kinase 2 | P54674 | 3E-63 | 68 | down |
| *pumA* | Pumilio homolog 2 | Q80U58 | 2E-46 | 80 | down |
| *anxA* | Annexin-B12 | P26256 | 6E-41 | 98 | down |
| *pptA* | Phosphatase DCR2 | Q05924 | 6E-19 | 59 | up |
| *cdcA* | Cell division control protein 31 | P06704 | 6E-27 | 38 | up |
| *pakA* | Serine/threonine-protein kinase pakC | Q55GV3 | 3E-48 | 79 | up |
| *pldA* | Phosphatidylinositol-glycan-specific phospholipase D | Q8R2H5 | 4E-62 | 91 | up |
| *ligA* | Checkpoint protein hus1 homolog 1 (LigA) | Q54NC0 | 1E-28 | 94 | up |
| *pwiA* | Piwi-like protein 1 | Q96J94 | 2E-55 | 92 | up |
| *rgsA* | Regulator of G-protein signaling 2 | O08849 | 3E-05 | 31 | up |

### 2.3. Statistical methods

The datasets for PHO3, for PHO57, and for the PHO3 + PHO57 heterokaryon were processed separately.

#### 2.3.1. Normalization of the gene expression data

Gene expression data were normalized to the values measured in the dark control time series. To do so, the mean of the expression values of all samples taken from cells that had not been exposed to a far-red light stimulus was calculated for each gene separately and the expression values for each gene measured in samples of far-red light stimulated plasmodia were normalized to the mean of the respective gene in the dark control time series.

#### 2.3.2. Generation of heat maps and hierarchical clustering

Clustering and visualization of the normalized gene expression data were performed with the function *heatmap.2* in R (Version 3.1.3). In parallel, significant cluster analysis (*α* = 0.001) was performed using the function *simprof* as part of the package clustsig [28]. Both functions employ the function hclust [29].

#### 2.3.3. Multidimensional scaling

After normalization of the gene expression data (see above), a structure representing the euklidian distance between the data points was computed by the function *dist* which is part of the *stats* library. Classical multidimensional scaling (MDS) of the gene expression data in two dimensions (*k*=2) was then performed on the distances using the function *cmdscale()* in R (Version 3.1.3) which is also part of the *stats* library and plotted with *plot()* contained in *graphics*.

## 3 Results

### 3.1. Light responsiveness and sporulation propensity of plasmodial cells is genotype-dependent

We analysed two strains, PHO3 and PHO57 that were isolated in a genetic screen for sporulation mutants [26]. As compared to the wild type, PHO3 displays a reduced propensity to sporulate in response to a far-red (FR) light stimulus while plasmodia of PHO57 do not sporulate at all in response to FR light stimulation. When starved PHO3 plasmodia were irradiated with a saturating pulse of far-red light, only approximately half of the population sporulated indicating that the probability for entering the developmental pathway is reduced as compared to the wild-type. However, the entire protoplasmic mass of each plasmodium sporulated as a whole or did not sporulate at all, indicating that the decision for sporulation to occur was still all-or-none in each plasmodium. Investigating the PHO3 mutant hence allows to study the dynamical bifurcation between the routes that lead or lead not to sporulation in the context of this genotype.

Plasmodial cells were starved for six days and exposed to a far-red light stimulus. Before and at different time points after the stimulus, a sample was taken from each plasmodium to analyse the expression pattern of a set of genes (Table 1) that are up- or down-regulated in the wild-type in association with developmental switching [15, 27] (Fig 1B). The remainder of the plasmodium was incubated until the next day to see whether sporulation had occurred. For the dark controls, plasmodia were not exposed to far-red light but otherwise treated identical. Comparative display of the time-series data obtained for each plasmodial cell of PHO3 revealed that a set of genes were up-regulated in the subpopulation of cells committed to sporulation as it had previously been found for the wild-type [27]. In order to group cells with respect to the similarity of the expression patterns, we pooled and normalized the data to obtain one dataset for the cells of each genotype (see Materials and Methods for details), performed a hierarchical cluster analysis of the expression values [29], and displayed the results in the form of a heat map (Fig. 2A).

**Figure 2.**
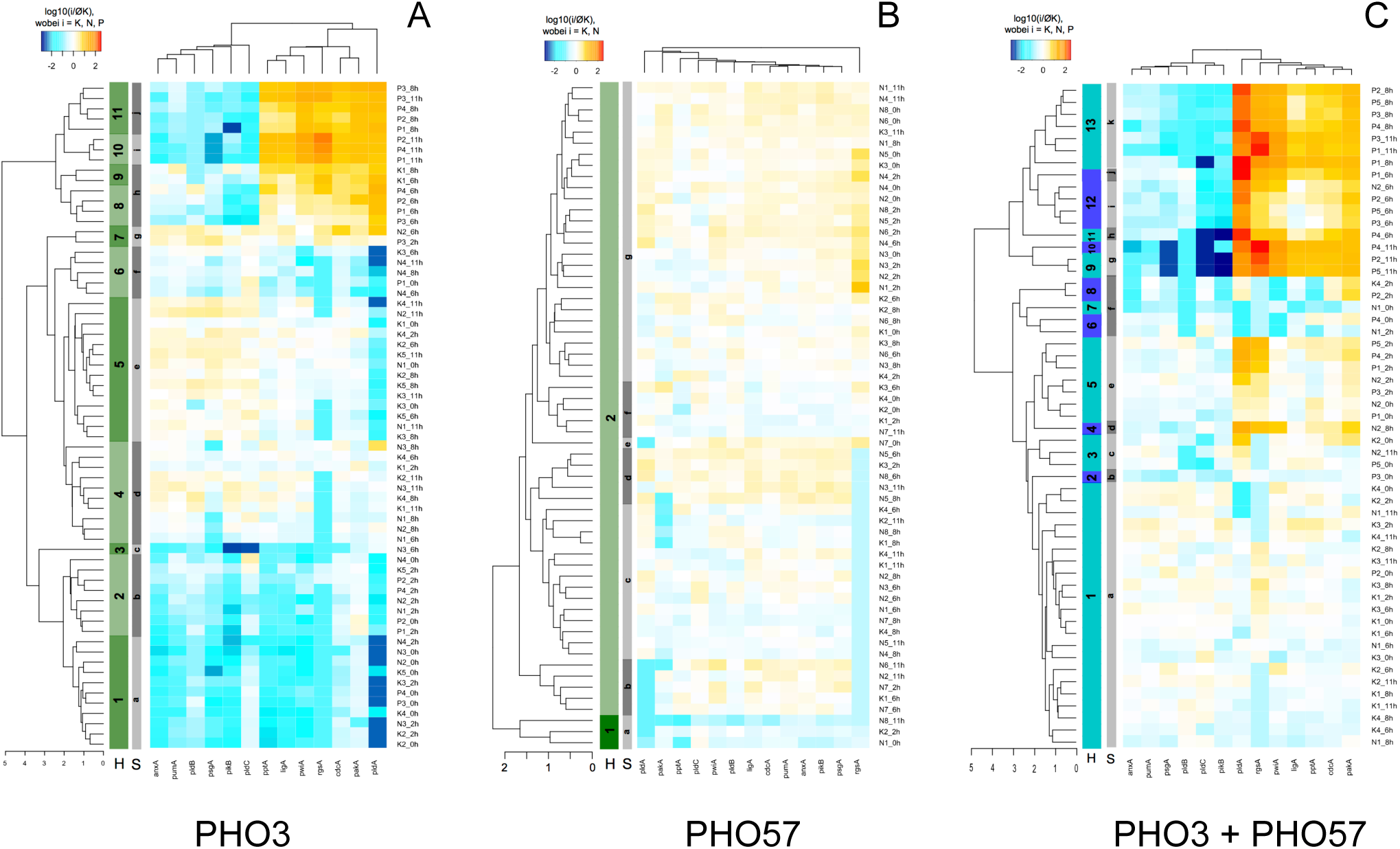
Differential gene expression patterns for a set of cell differentiation marker genes in response to far-red light stimulation of plasmodial cells. Heat maps and clusters as obtained by hierarchical clustering (H) or simprof analysis (S) are shown for of PHO3 (A), PHO57 (B) and for the PHO3 + PHO57 heterokaryon (C). Time series were obtained by repeatedly taking samples from plasmodial cells that were stimulated with far-red light or that were not stimulated (dark control time series), giving one time series for each individual plasmodial cell together with the information whether or not the remainder of the plasmodium had sporulated (Fig. 1B). The data points of three groups of plasmodial cells (unstimulated, not sporulated; stimulated, not sporulated; stimulated and sporulated) were pooled and the three resulting datasets (for PHO3, PHO57 and for the PHO3 + PHO57 heterokaryon) used for hierarchical clustering and the heat maps (this Figure), for multidimensional scaling (Fig. 3), and for the reconstruction of the single cell trajectories (Fig. 4).

PHO57 cells did not sporulate in response to a far-red light stimulus which is saturating in wild type [26], and there was no obvious response in differential gene expression (Fig. 2B).

Heterokaryons of equal relative cytoplasmic mass of PHO57 and PHO3 were obtained by fusing plasmodial cells of both strains. Accordingly, the PHO3 and PHO57 mutant gene dosages were approximately 50 % in the heterokaryon while the presence of a dosage of 50 % of the corresponding wild-type alleles of the respective fusion partner may cause complementation. The heterokaryons showed a qualitatively similar phenotype as PHO3 cells. As in PHO3, only approximately 50 % of the plasmodial cells of the PHO57 + PHO3 heterokaryon population sporulated in response to a saturating far-red light stimulus, suggesting that the PHO57 mutation in conjunction with PHO3 is recessive at a gene dosage of 50 %. Sporulated cells qualitatively showed a similar response at the gene expression level (Fig. 2C) as PHO3 cells did, i.e. down-regulation of *psgA*, *pldC*, *pldB*, *pikB*, *pumA*, *anxA* and up-regulation of *pptA*, *cdcA*, *pakA*, *pldA*, *ligA*, *pwiA*, and *rgsA* (Table 1).

### 3.2. Multidimensional scaling analysis reveals variable trajectories of individual cells

To investigate the variability of pathways a plasmodium can take to commitment and sporulation and the dependence of these pathways on the genotype, we performed a multidimensional scaling (MDS) analysis [30] to arrange the data points according to their similarity as projected into a two-dimensional plane. Clearly, data points were unevenly spread over the area of the plot and there were certain regions where data points from sporulated or from non-sporulated plasmodia were predominantly located (Fig. 3; red *vs*. black or blue data points, respectively). Data points that were assigned to a cluster by hierarchical clustering (Fig. 2) mapped to a corresponding cloud of points in the MDS plot (Fig. 3; dashed lines), indicating that the obtained results were consistent. MDS analysis of the gene expression data of PHO57 cells revealed data points in one significant cluster with three outliers that mapped to a second cluster, suggesting that the far-red light exposure did not induce any significant response in gene expression. We then reconstructed the trajectory for each individual cell by connecting the data points in the MDS plots according to the temporal order in which the samples had been taken from the cell. The trajectories of the sporulated plasmodia (indicated in red in Fig. 4) moved towards the lower part of the MDS plot as compared to the trajectories of the non-sporulated plasmodia of PHO3 and of the PHO3 + PHO57 heterokaryon (Fig. 4A,C). One feature however was the same for all plasmodia of PHO3 and of the heterokaryon. Their trajectories spanned regions corresponding to more than one cluster, suggesting that plasmodia switched between different gene expression patterns, even those cells that had not been stimulated by a light pulse. Corresponding analyses performed with the non-sporulating strain PHO57 (Fig. 4B) and the differences between PHO3 and the PHO3 + PHO57 heterokaryon suggest that cluster formation, MDS patterns, and single cell trajectories depend on the genotype that influences the propensity of a plasmodium to sporulate (see below).

**Figure 3.**
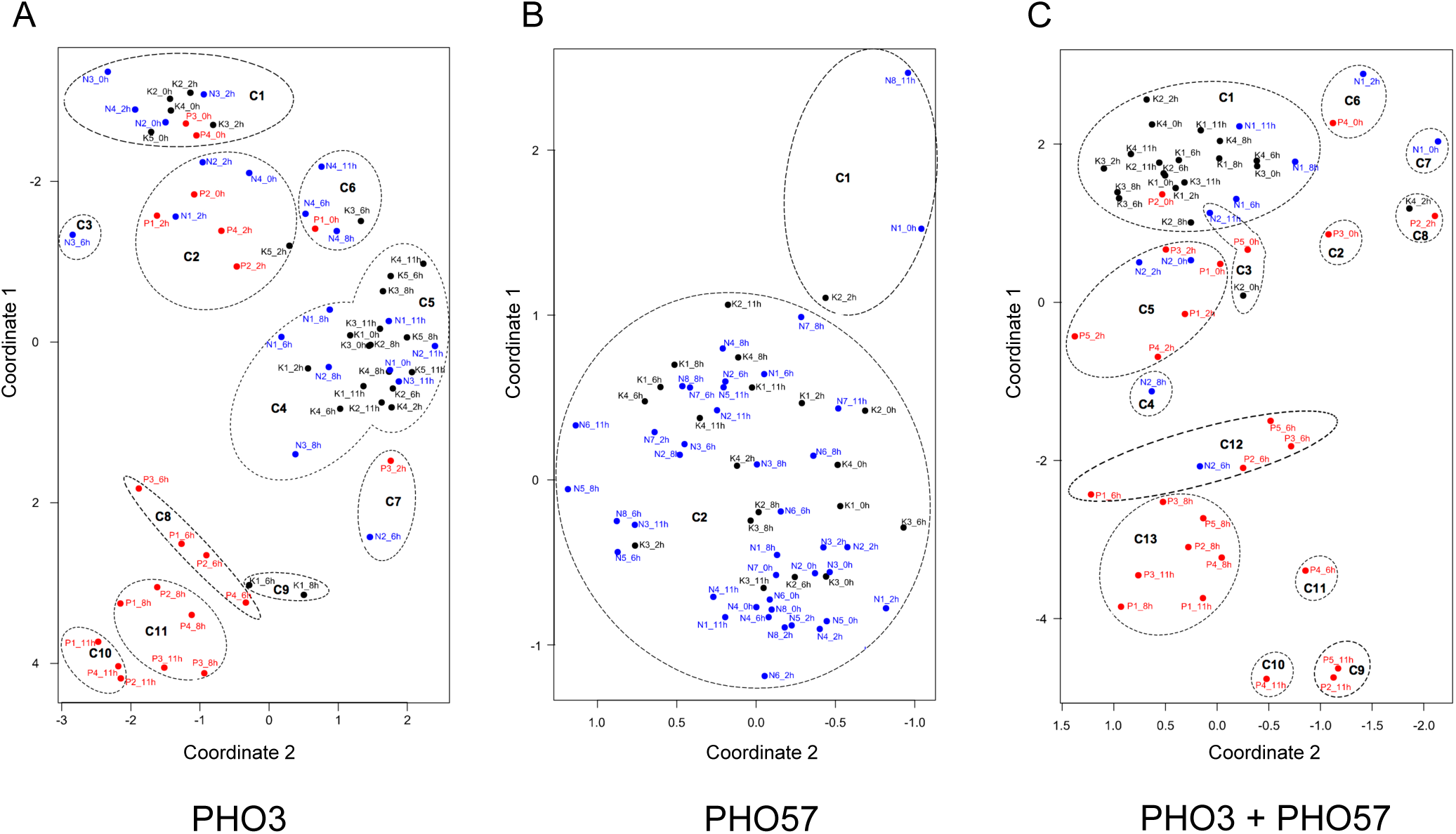
Multidimensional scaling (MDS) of the data points of cells of PHO3 (A), PHO57 (B) and of the PHO3 + PHO57 heterokaryon(C). Each data point represents the gene expression pattern of a cell at a given time point. The data points in the MDS plot were assigned to a respective cluster found by hierarchical clustering in (Fig. 2) and were accordingly marked with a dashed line.

**Figure 4.**
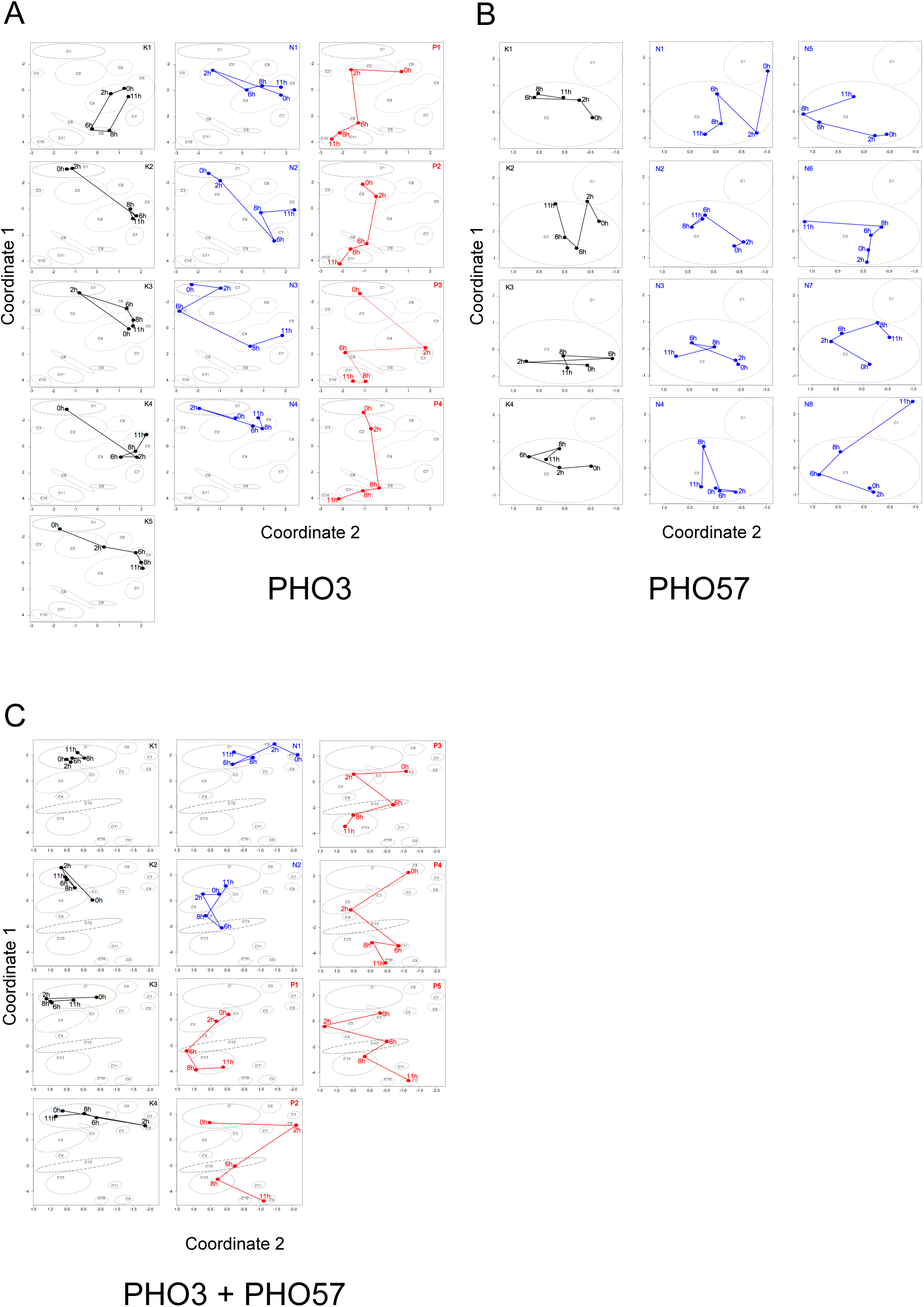
Single cell trajectories of differential gene expression. Data points of the time series from the multidimensional scaling plot of (Fig. 3) were connected to give a trajectory representing the gene expression patterns of subsequent time points for each individual plasmodial cell. Spatial regions as assigned to the clusters as determined in (Fig. 2) were outlined as in (Fig. 3). Color coding of trajectories: not stimulated, not sporulated, black; far-red light stimulated, not sporulated, blue; far-red light stimulated, sporulated, red.

### 3.3. Reconstructing a Petri net model of state transitions from single cell trajectories

The two methods, hierarchical clustering and MDS gave consistent yet complementing results. While MDS visualizes the similarity of the expression patterns of samples in a two-dimensional plane, clustering assigns samples to groups resulting in a statistically significant discretization. Based on this discretization, we reconstructed a Petri net model from the trajectories of individual cells of each strain, PHO3, PHO57, and of the PHO3 + PHO57 heterokaryon. The gene expression pattern of each plasmodium at each time point as defined by its assigned cluster was listed in a table to obtain a comprehensive representation of all time series (Table 2). This table was translated into a Petri net that models the transitions between the gene expression states of each strain. For a brief introduction to the Petri net formalism including the elements used in this study, see Box 1 and Fig. 5.

**Table 2.** Single cell trajectories of gene expression patterns as discretized by hierarchical clustering. Each line represents the time series measured in an individual plasmodial cell. Cells were labled as follows. K, not stimulated, not sporulated, N, far-red light stimulated, not sporulated, P, far-red light stimulated, sporulated. Clusters (H) are marked in the heat maps of Fig. 2. Clusters that were found in the cells of the dark controls are shown in black, those that were found in light stimulated but not in sporulating cells in blue, and those that were exclusively found in sporulating cells are shown in red.

Cells of strain PHO3
| Cell \ time | 0h | 2h | 6h | 8h | 11h |
| --- | --- | --- | --- | --- | --- |
| K1 | C5 | C4 | C9 | C9 | C4 |
| K2 | C1 | C1 | C5 | C5 | C4 |
| K3 | C5 | C1 | C6 | C5 | C5 |
| K4 | C1 | C5 | C4 | C4 | C5 |
| K5 | C1 | C2 | C5 | C5 | C5 |
| N1 | C5 | C2 | C4 | C4 | C5 |
| N2 | C1 | C2 | C7 | C4 | C5 |
| N3 | C1 | C1 | C3 | C4 | C4 |
| N4 | C2 | C1 | C6 | C6 | C6 |
| P1 | C6 | C2 | C8 | C11 | C10 |
| P2 | C2 | C2 | C8 | C11 | C10 |
| P3 | C1 | C7 | C8 | C11 | C11 |
| P4 | C1 | C2 | C8 | C11 | C10 |

Cells of strain PHO57
| Cell \ time | 0h | 2h | 6h | 8h | 11h |
| --- | --- | --- | --- | --- | --- |
| K1 | C2 | C2 | C2 | C2 | C2 |
| K2 | C2 | C1 | C2 | C2 | C2 |
| K3 | C2 | C2 | C2 | C2 | C2 |
| K4 | C2 | C2 | C2 | C2 | C2 |
| N1 | C1 | C2 | C2 | C2 | C2 |
| N2 | C2 | C2 | C2 | C2 | C2 |
| N3 | C2 | C2 | C2 | C2 | C2 |
| N4 | C2 | C2 | C2 | C2 | C2 |
| N5 | C2 | C2 | C2 | C2 | C2 |
| N6 | C2 | C2 | C2 | C2 | C2 |
| N7 | C2 | C2 | C2 | C2 | C2 |
| N8 | C2 | C2 | C2 | C2 | C1 |

**Cells of the PHO3 + PHO57 heterokaryon**
| Cell \ time | 0h | 2h | 6h | 8h | 11h |
| --- | --- | --- | --- | --- | --- |
| K1 | C1 | C1 | C1 | C1 | C1 |
| K2 | C3 | C1 | C1 | C1 | C1 |
| K3 | C1 | C1 | C1 | C1 | C1 |
| K4 | C1 | C8 | C1 | C1 | C1 |
| N1 | C7 | C6 | C1 | C1 | C1 |
| N2 | C5 | C5 | C12 | C4 | C3 |
| P1 | C5 | C5 | C12 | C13 | C13 |
| P2 | C1 | C8 | C12 | C13 | C9 |
| P3 | C2 | C5 | C12 | C13 | C13 |
| P4 | C6 | C5 | C11 | C13 | C10 |
| P5 | C3 | C5 | C12 | C13 | C9 |

**Figure 5.**
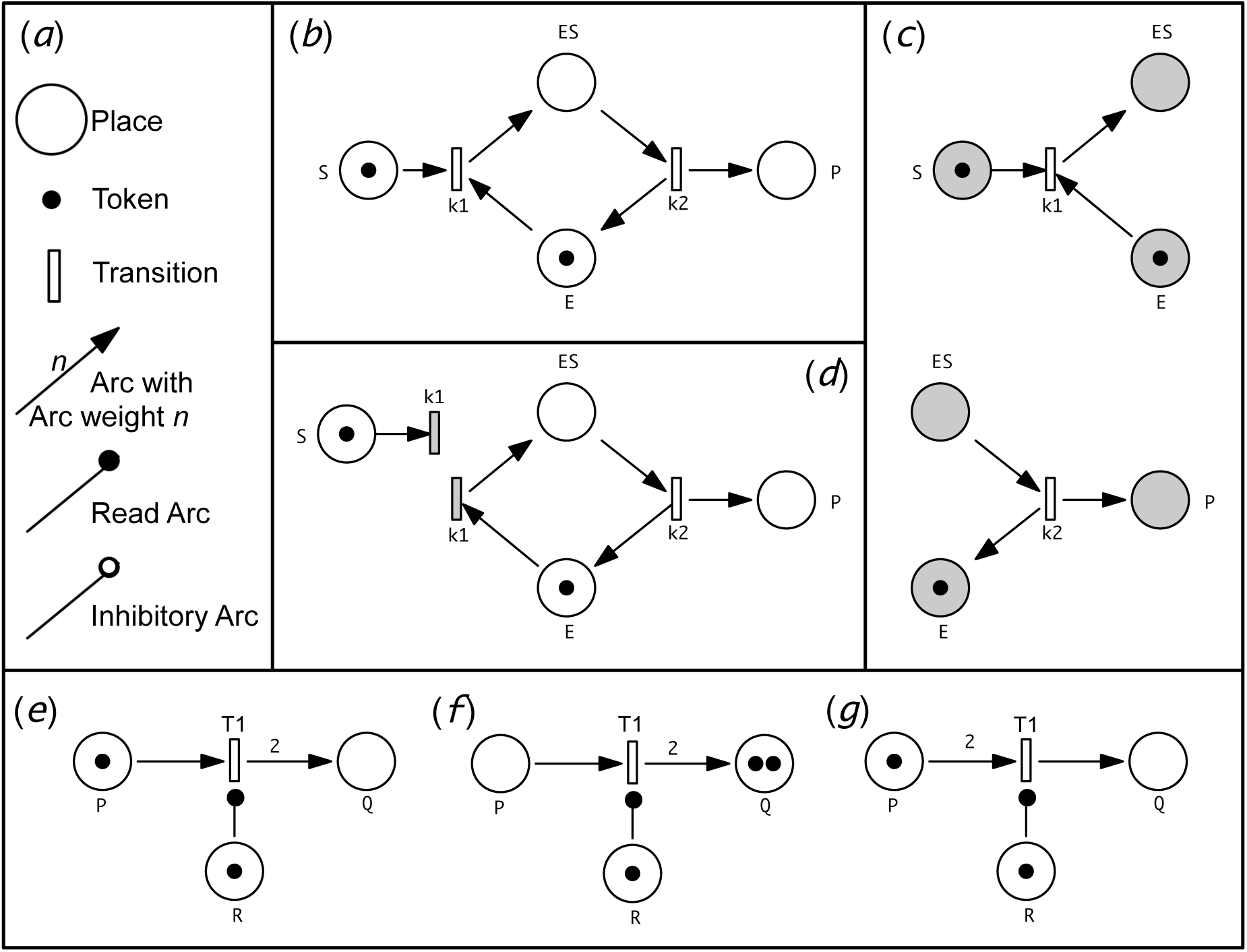
Elements of extended Petri nets used in this study and their functionality. (a) Petri nets are bipartite directed graphs in which two types of nodes, places and transitions, are mutually connected by arcs. Extended Petri nets additionally provide read arcs and inhibitory arcs which link a place to a transition to control the enabledness of the transition. (b) Petri net representing an enzyme-catalyzed biochemical reaction in the form of E+S → ES → E+P, for simplicity neglecting the reversibility of the reactions. In the example, the places representing substrate and enzyme, respectively, carry one token each, meaning that there is one molecule of substrate and one molecule of enzyme. Transitions represent biochemical reactions. Upon firing, transition k_1_ takes one token from S and one token from E and delivers one token in the place representing the enzyme-substrate-complex. As soon as there is a token in place ES, transition k_2_ is enabled and may consume the token from ES to put one token each in places P and E. Subsequently, none of the transitions can fire any more, as there is no token in S. In stochastic Petri nets, firing of a transition occurs with a constant probability per unit of time as defined by a rate constant as soon as the transition is enabled. (c), (d) The graphical copy of a node is called logic node and can be used to graphically disentangle a Petri net. In the Petri net tool Snoopy, a node is automatically shaded if it is defined as logic. (c) The Petri net of (b) is represented using logic places for all reactants S, E, ES, and P. Note that the tokens by which a logic place is marked are automatically displayed in all graphical copies of this place. (d) The Petri net of (b) with k_1_ defined as a logic transition. Accordingly, k_1_ can only fire if the two places, S and E, contain at least one token. (e,f,g) Use of read arcs and the meaning of arc weights. The arc weight is indicated as a number next to an arc. If no arc weight is given, the arc weight by definition is one. As place R is connected to transition T_1_ with a read arc with an arc weight of one, the transition can only fire if R is marked by at least one token. The arc weight of standard arcs corresponds to the stochiometry of token displacement. In other words, it indicates the number of tokens that are transported via the arc connected to the transition when this transition fires. If a pre-place of a transition contains a number of tokens which is less than the arc weight of a standard arc, or a read arc which is directed to the transition, the transition cannot fire. The marking of a place that is connected via a read arc or an inhibitory arc to a transition does not change when the transition fires. Simply said, read arcs and inhibitory arcs sense rather than transport tokens. When T_1_ in the Petri net of (e) fires, it takes, according to the arc weights, one token from P and puts two tokens into Q while the marking or R remains unchanged (f). Transition T_1_ in the Petri net of panel (g) cannot fire because there is only one token in P while the arc weight requires that two tokens are consumed from P upon firing of the transition. If place R is connected to transition T_1_ with an inhibitory arc instead of the readarc in (e), T_1_ can only fire if there are less tokens in place R than indicated by the arc weight of the inhibitory arc (not shown). A brief introduction to Petri nets and to the Petri net tool Snoopy can be found in [37].

For creating a Petri net model that is able to reproduce and predict single cell trajectories by simulation, a place was assigned to each cluster as defined by hierarchical clustering (Table 2) and MDS. For the initial graphical representation, the places were put in a relative geometric position that corresponds to the center of each cluster as it is located in the MDS plot (Fig. 3). The places were then connected by transitions in a way that each single cell trajectory could be represented by the token moving through the Petri net. The token corresponds to the marble in Waddingtons metaphor (see Discussion). To graphically discriminate the pathways as they were observed in a particular group of plasmodial cells, arcs directing the pathways taken by the dark controls were indicated in black, those of far-red light irradiated but not sporulated cells in blue and those of far-red light-induced and sporulated cells in red (Fig. 6). Paths that were taken by plasmodia from more than one group were encoded with arcs of two or more colors (e.g. arcs connecting transitions T12 or T16 in Fig. 6). Note that the coloring of the arcs is only for illustration or visualisation purposes but does not have any functional relevance for the Petri net. Note also, that this is not a "colored Petri net" in the technical sense. Colored Petri net is a technical term for a special class of Petri nets [38].

**Figure 6.**
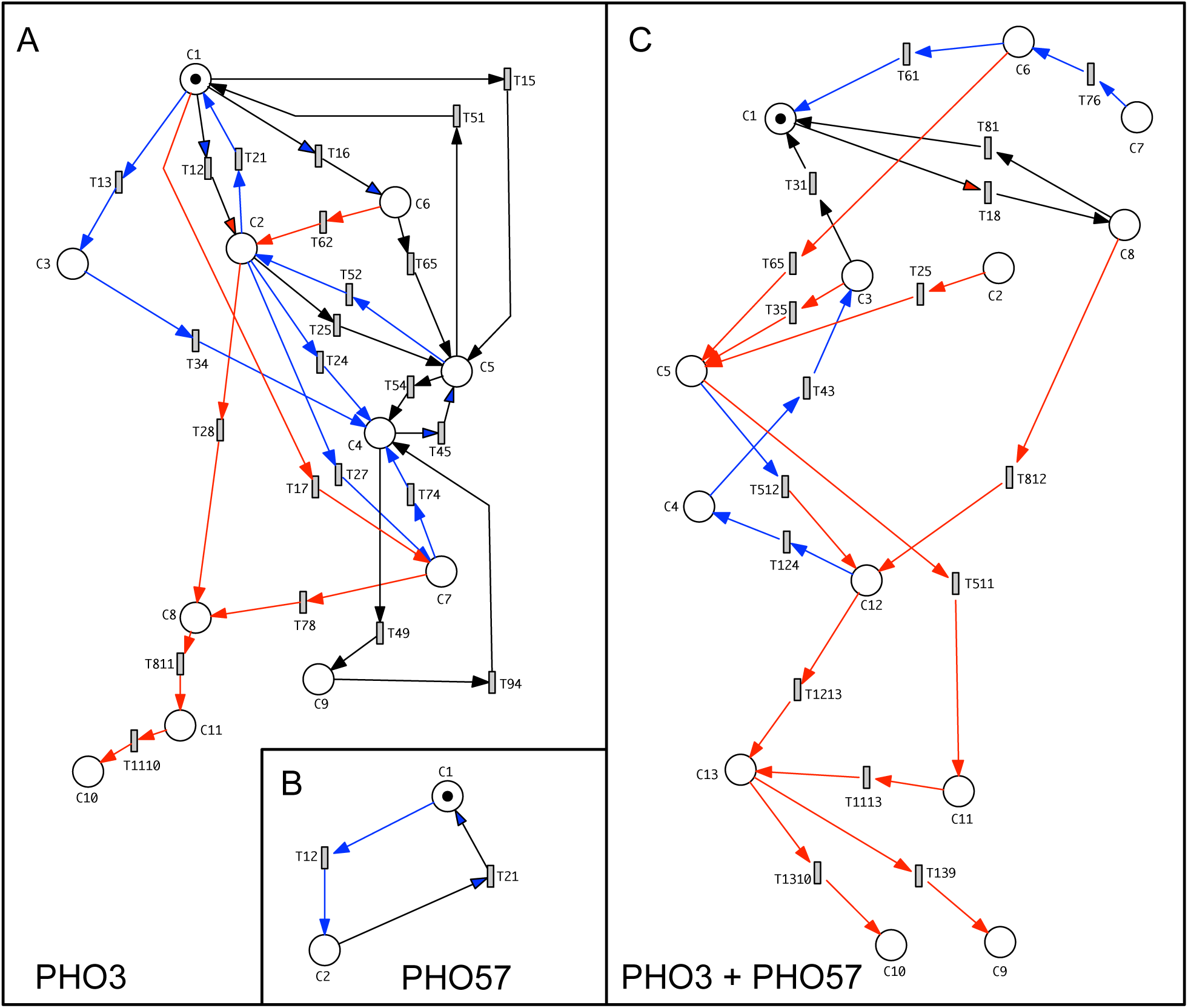
Petri nets reconstructed from the time series of transitions between states of gene expression of PHO3 (A), PHO57 (B) and of PHO3 + PHO57 heterokaryon (C) cells. Places representing the clusters as determined in (Fig. 2) and listed in Table 2 were geometrically arranged relative to each other corresponding to the center of each cluster in the MDS plot of Fig. 3 so that the information regarding the similarity of expression patterns between the clusters is encoded in the relative spatial position of the Petri net places. Transitions of the Petri net were named according to the places they connect, e.g. transition T12 connects place C1 to place C2 through directed arcs. The color coding of the arcs indicates the group of cells in which the paths were observed: not stimulated, not sporulated, black; far-red light stimulated, not sporulated, blue; far-red light stimulated, sporulated, red. The Petri net place representing the starting point of a single cell trajectory is marked with one token. Upon firing of enabled transitions, the token indicating the gene expression state of the cell moves through the net and creates a trajectory similar to the trajectories in (Fig. 4).

##### Box 1. Standard and extended Petri nets

Petri nets are mathematical structures that are broadly used to model concurrent systems with classical applications in computer science, systems engineering, and operations research [31]. Petri nets are also common to model, analyse, and simulate biochemical reaction networks [32, 33]. Petri nets are bipartite directed graphs that are composed of places, transitions and directed arcs (Fig. 5). In biochemical reaction networks, places may represent biochemical components while transitions refer to biochemical reactions [33]. In a more general sense, a place may represent an entity or a certain state. When a place contains one or more tokens this means that the entity is present in the copy number as indicated by the number of tokens in the place. To indicate whether a state is true, one token is sufficient. A transition in a Petri net is always connected to one or more places by at least one directed arc. Upon firing, the transition can transport a token in the direction defined by the connecting arc and thereby change the marking of the connected places (for details see Fig. 5). By firing of a transition and moving of at least one token, the system accordingly transits from one state into another. Petri nets can be used as a formal language that is compatible with virtually all modeling paradigms common in systems biology [34, 35], including Boolean netoworks. Petri nets have been employed to model the transition between physiological states and genetic complementation effects in the photosensory control of sporulation in *Physarum polycephalum* [36].

In Fig. 6, the coloring of the arcs highlights that there were pathways that were exclusively taken by those plasmodia that had sporulated and some that were predominantly taken by light-exposed plasmodia that did not sporulate, while others were perdominantly taken by the dark controls. Certainly, the low number of plasmodial cells in each sample does not allow for any final conclusion regarding the exclusiveness of the pathways which are taken in response to a certain treatment. Nevertheless, our case studies demonstrate how a Petri net analysis of single cell trajectories on MDS data can be performed. With the Petri net, single cell trajectories can be simulated as follows: One token is put into a place defining the state in which the plasmodium is when the trajectory starts. The token is then allowed to move spontaneously through the stochastic Petri net as enabled transitions fire randomly giving a simulated single cell trajectory for each simulation run. Simulations can be performed with Snoopy [35, 39], the tool that was used to draw the Petri nets. The relative frequency of a transition to fire and hence the relative frequency of individual trajectories to occur results from the firing propensity of the transitions (corresponding to kinetic rate constants) that can be defined in the stochastic Petri net. Even without running a simulation, the Petri net visualizes the pathways that were taken by individual cells on their walk through the quasipotential landscape of gene expression (see Discussion) and it displays structural properties that are analysed in the following.

### 3.4. T-invariant analysis indicates cyclic transition between states

Before we assign the firing of the transitions to the differential regulation of specific genes, let us first consider some structural properties of the Petri net. We discuss the properties of the nets in their current form, bearing in mind that the reconstruction is based on a limited amount of data. With more cells analysed, the nets might well become more structured with more places and more transitions. Also, currently observed differences in trajectories between groups of cells (e.g. light-exposed *vs.* dark controls) might emerge or disappar with more cells analysed or with more time points taken. Despite of this caveat, the currently available dataset allows to demonstrate the computational method.

The places in the Petri net of Fig. 6 have been purposefully arranged to visually assign them to the relative positions of the clusters within the MDS plot. In this format, the relative position of the places indicates how similar the clusters are relative to each other with respect to the gene expression patterns they comprise. In order to arrange the nets so that their structure appears more clearly, we used an automatic layout algorithm implemented in Snoopy (Fig. 7). Although the connectivity within the re-arranged nets is unchanged, the information regarding the similarity of the clusters is lost as the relative positions of the places corresponding to the relative positions of the clusters in the MDS plot is not preserved. The rearranged net was then used to analyse the T-invariants. A T-invariant can be described as follows: If each transition belonging to a T-invariant in partial order has fired once, the marking of the places of the T-invariant is the same as it was before the first transition of the T-invariant fired (see color-coded T-invariants in Fig. 8 as examples; for a formal definition of T-invariants see [40]. In other words, a T-invariant defines a reaction cycle or a sequence of firing events that brings the subsystem which is covered by this T-invariant back to its original state.

**Figure 7.**
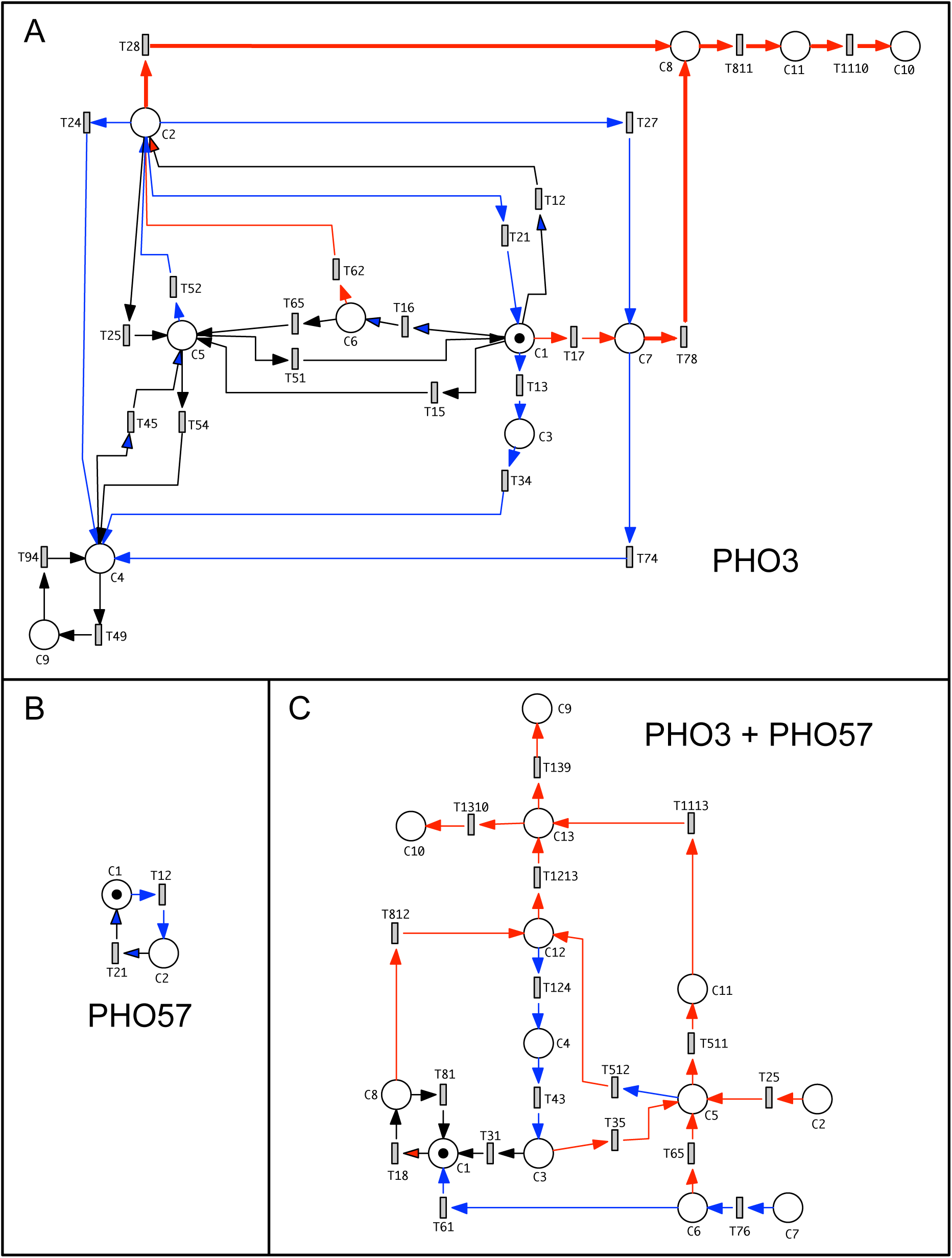
Planarized Petri nets used for the analysis of T-invariants. The Petri nets of Fig. 6 were subjected to an automatic layout algorithm (planarization) implemented in the Petri net tool Snoopy in order to get the graphical representation clearly arranged. The new layouts were used to visualize the T-invariants as calculated with the analysis tool Charlie [41]. The color coding of the arcs in (A), (B), and (C) indicates in which types of plasmodial cells the paths were observed: Dark controls, not sporulated, black; far-red stimulated, not sporulated, blue; far-red stimulated, sporulated, red. The Petri net of PHO3 cells (A) contains 21 extensively overlapping T-invariants which are shown in Supplemental Movie 1. The simple net of PHO57 (B) consists of one single T-invariant. Transitions in (A) were defined as logic transitions to allow the connections to those parts of the model that were added to represent light regulation, differential gene expression, and the physiological states of the cell (Fig. 9). The three T-invariants of the net of the PHO3 + PHO57 heterokaryon (C) are shown in Fig. 8.

**Figure 8.**
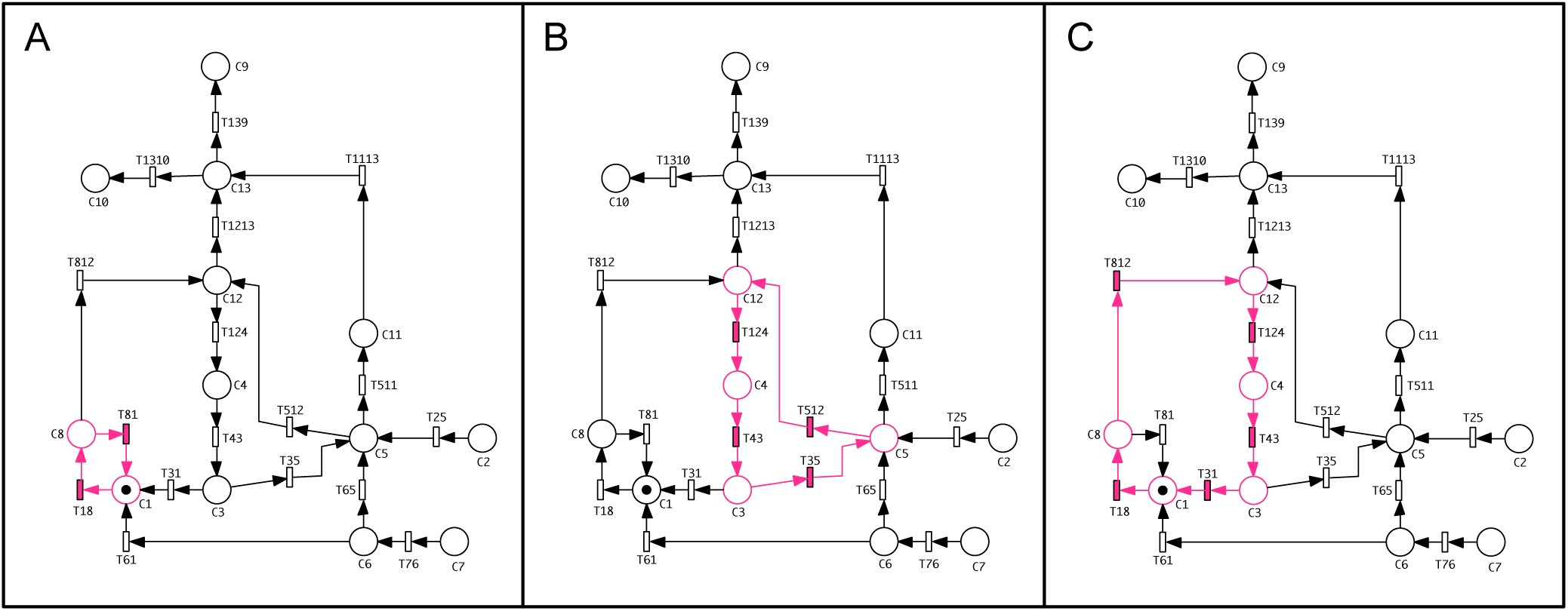
T-invariants of the Petri net modeling the transitions between changes between states of gene expression in the PHO3 + PHO57 heterokaryon. The T-invariants were computed and displayed using the analysis tool Charlie [41] and displayed in Snoopy as shown in Supplemental Movie 1.

T-invariants were calculated using the analysis tool Charlie [41] and subsequently visualized in Snoopy. The net of PHO3 cells in Fig. 7A is mostly covered by a set of 21 T-invariants that display multiple overlaps. Overlapping is so extensive that color-coding of all individual T-invariants in the net of Fig. 7A is not even possible. As inspection of the individual T-invariants is instructive, the calculation of T-invariants with Charlie and visualization of the complete set with Snoopy is shown in the Supplemental Movie SI Movie 1. T-invariants only occurred in those parts of the net that were reconstructed from the trajectories of cells that did not sporulate. Therefore, the red bold arcs in Fig. 7A highlight those regions of the net that are not part of any of the 21 T-invariants and that were taken by the sporulated cells only. T-invariant analysis reflects that non-sporulated cells cyclically switchend between different states of gene expression. It further predicts numerous trajectories that would emerge from the combinatorics of state transitions. From the color-encoded arcs in Fig. 7A it seems that far-red light exposure created T-invariants in addition to those emerging from the trajectories of the dark controls. However, the low number of cells analysed for each group does not allow for any final conclusion. With this limitation in mind, the T-invariants involving C3 or C7 might well be stimulus-specific Fig. 7A. T13, the transition from C1 to C3 (Fig. 7A) with a strong differential regulation of *pikB, pldC* (down), and *pldA* (up) may indicate a futil attempt of PHO3 cells to switch to the cell fate of sporulation.

In summary, the high number of T-invariants reflects that non-sporulated PHO3 cells were switching between states of gene expression. Cells that sporulated in response to the far-red light stimulus irreversibly proceeded to clusters C8, C11 and C10, however on seemingly different pathways C2 to C8 or C1 to C8 via C7 according to the Petri net and as suggested by the experimental results shown in Table 2. Based on the available dataset we cannot exclude that C7 represents a state with short life time that might also have occurred upon the transition from C2 to C8 though in might not have been resolved due to the low number of time points that were sampled. Anyway, the high number of T-invariants observed for PHO3 cells was not found in the PHO3 + PHO57 heterokaryon (Fig. 8) indicating that the structure of the Petri net reflecting the topology of the quasipotential landscape is genotype-dependent (see Discussion).

The trajectories of PHO3 cells that were committed to sporulation moved through cluster C8 to clusters C11 and eventually to C10. In any temporal sequence of events, C8 was the first place which was visited by all cells that subsequently sporulated. Only cells that were committed to sporulation entered C8, suggesting that transitions T28 and T78 represent the committment to sporulation (Fig. 7A) where the system is trapped by a new basin of attraction (see Discussion).

The Petri net obtained for PHO57 cells (Fig. 7B) is accordingly simple as there were only two significantly different clusters in the MDS plot. The net consists of a single T-invariant.

The Petri net reconstructed from the pathways of cells of the heterokaryon Fig. 7C was different as compared to the Petri net for the PHO3 cells with its 21 T-invariants. In the Petri net for the heterokaryon there were only three T-invariants, one for the cells of the dark controls and two for light-exposed cells (Fig. 7C; Fig. 8). The other pathways lead to sporulation. There were two nodes of pre-activation for sporulating cells, C5 and C12 from which cells could return without sporulating. And there was one central node C13 for all sporulating cells which characterizes a state at which the cells had been committed to sporulation. The difference in the number of T-invariants indicates that the behaviour of the cells of the PHO3 + PHO57 heterokaryon is more directed than the behaviour of the PHO3 cells suggesting that the topology of the quasipotential landscape is genotype-dependent. This more directed behaviour with less T-invariants within the Petri net may well be due to partial complementation of the sporulation-suppressing PHO3 mutation by the wild type gene product contributed by the PHO57 plasmodium in the heterokaryon.

### 3.5. Light-dependent steps and commitment of plasmodial cells

Single cell trajectories have been obtained for three groups of cells, for far-red light-exposed cells that sporulated or that did not sporulate in response to the stimulus and for dark controls that did not sporulate either. Arcs representing trajectories taken by irradiated, not sporulating cells only are indicated in blue in Fig. 7A-C. However, due to the low number of samples it cannot be excluded that dark control cells would not take at least certain steps of these pathways. Hence we searched for the maximal number of transitions that according to the net structure would be expected to be light-dependent. To do so, we identified those significant clusters that did not occur in cells of the dark controls and instead were found in light-stimulated cells that did not sporulate (marked in blue in Table 2). In addition, we identified those clusters that exclusively occurred in cells that sporulated in response to light stimulation (marked in red in Table 2). With the caveats mentioned above, transitions leading to places representing these clusters are considered to be directly or indirectly light-dependent. For PHO3 cells these are transitions T28 which, as discussed above, is directly associated with commitment to sporulation and T13, T17, and T27 that lead to C3 or to C7. Accordingly, C3 and C7 are candidates for states of pre-activation or induction, respectively which might either decay via transitions T34 or T74 or lead to commitment via T78. Based on these considerations, the minimal model for light activation of PHO3 cells is shown in Fig. 9. The phytochrome photoreceptor is activated by photoconversion of P_fr_ by far-red light. Photoconversion then enables the logic transitions T13, T17, T27, and T28, meaning that the three transitions cannot fire without any far-red light stimulus. Coupled through the so-called logic transitions, highlighted in grey, the light-activation module of Fig. 9A is a functional part of the Petri net of Fig. 7A. Graphical copies of a place or a transition are called logic places or logic transitions. The principle is explained in Fig. 5c,d. The use of logic nodes (logic places and/or logic transitions) allows to stucture and to modularize the graphical appearence of the Petri net having one part that models the MDS results and simulates the single cell trajectories (Fig. 7A) and other functionally coupled but graphically separate modular parts that model molecular events, photoreceptor activation, or changes in the physiological state of the cell in response to e.g. the photoactivation of the phytochrome (Fig. 9).

**Figure 9.**
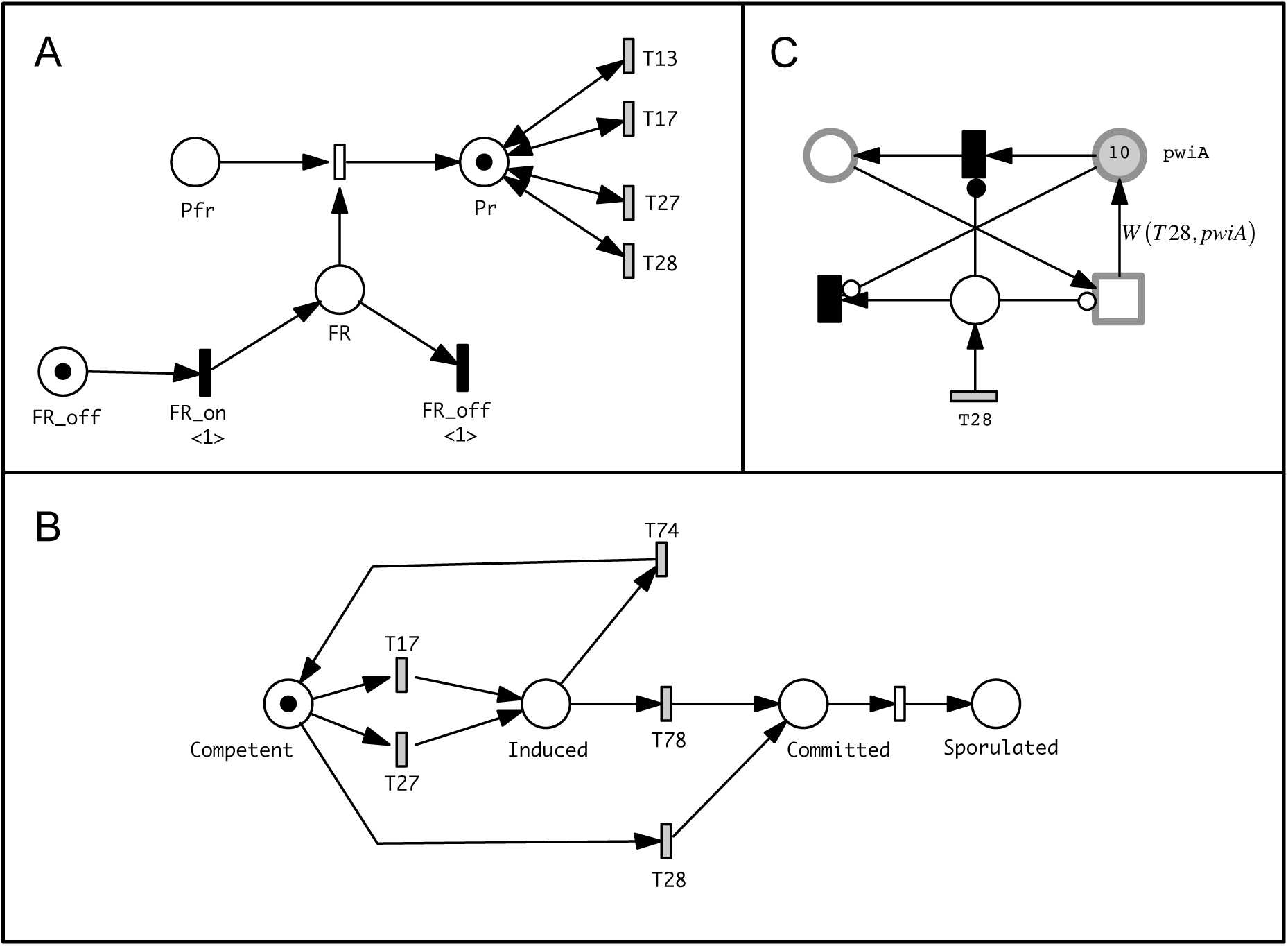
Petri net modules representing the phytochrome-mediated photosensory control of transitions between states of gene expression (A), associated changes in physiological states (B), and the differential expression of individual genes (C) in PHO3 cells. (A) Photosensory control. Places represent absence or presence of far-red light (FR) and the phytochrome photoreceptor in its P_fr_ and P_r_ states, respectively. Logic transitions are shaded in grey. Transitions filled in black are so-called immediate transitions that fire without delay when the preplace is sufficiently marked (for details see [37]). (B) Physiological states of the plasmodium, their photosensory control and their relation to the transitions between the clusters of differential gene expression as shown in Figs 2,3, and 7. (C) Sample copy of a gene expression module, updating the expression level of a gene upon firing of a transition linking the places representing clusters shown in (Fig. 7A). In order to model the differential regulation of all genes, this module is required in multiple copies (= number of transitions in Fig. 7A x number of genes represented). Its structure is designed to update the expression level of a gene according to its x-fold change entered as an arc weight. Those places and transitions that perform a computational function only (and do not have any biological interpretation) have not been labelled by a name. The network motif in the example shown for the *pwiA* gene is triggered by one of the transitions in (A), T28. As T28 fires, the expression level of the *pwiA* gene represented by the marking of the pwiA place is retrieved and updated according to the arc weight W(T28, pwiA). The arc weights for the set of analysed genes are displayed in Table 3 as x-fold changes in the gene expression level upon firing of transitions T28 and T78 which represent the commitment to sporulation. Supplementary Table 2 correspondingly displays the arc weights for all transitions of panel (A). Places representing the gene expression level are implemented and graphically displayed as continous places [42] and can therefore be updated in the form of continuous numbers. As the Petri net contains stochastic and continuous nodes, it is called hybrid Petri net. Note that the gene expression modules are not more than entities that, in the context of the coherent model, phenotypically mimic and predict trajectories of differential gene expression but they do not represent any regulatory mechanism at the molecular level.

In an analogous line of argumentation, transitions leading to commitment can be identified with the help of Table 2 and employed in a modular part of a Petri net to assign changes of the physiological state of a cell to the transition between clusters. In PHO3, transition to cluster C8 that was only seen in cells committed to sporulation occurs via T28 or T78 while induction is mediated via T17 or T27 (Fig. 9). In cells of the heterokaryon, T1213 and T1113 lead to commitment, while the state of induction decays via T124 (Table 2).

**Table 3.**
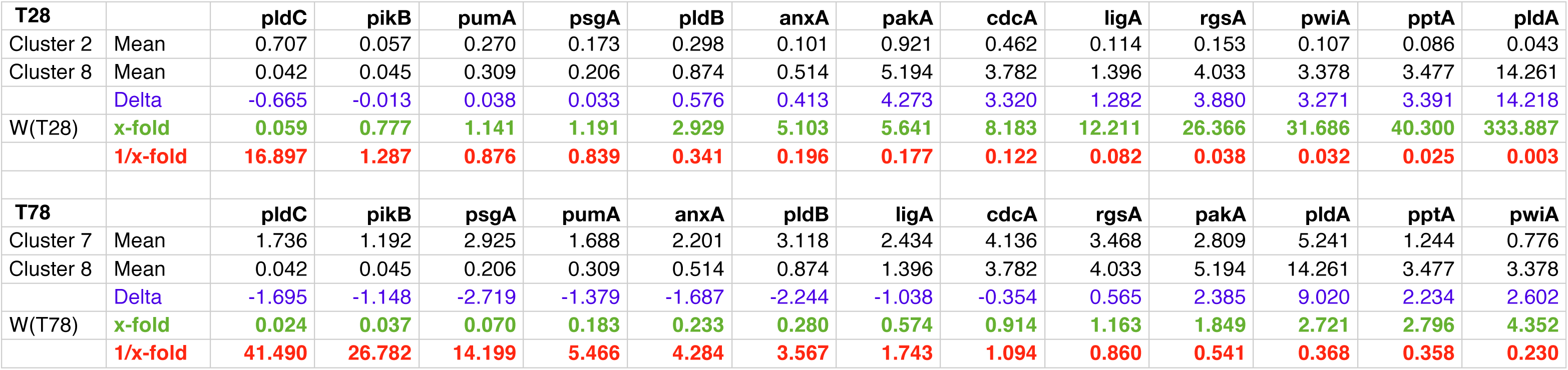
Differential gene expression associated with the commitment to sporulation of PHO3 cells. The x-fold changes in gene expression associated to firing of transitions T28 and T78 in the Petri net of Figs 6A,7A were calculated by taking the means of the expression values of each gene in the two clusters that were compared. The values for T28 and for T78 were then sorted for ascending x-fold changes. In the Petri net of Figs 6A,7A, the arc weight W(Tx,genX) entered into each gene expression module (Fig. 9C) corresponds to the x-fold change of each gene for the respective transition as it is provided by this table. The x-fold changes of gene expression values for all transitions of the Petri net of Figs 6A,7 is provided as Supplementary Table 2.

### 3.6. Formal assignment of the transition between clusters to the differential regulation of specific genes

The Petri nets in Figs. 6 and 7 model how the collective state of a cell may change with time depending on whether or not the cell has received a light stimulus. However, these Petri nets are solely referring to the states as defined by gene expression patterns discretized by clustering. In order to consider the differential regulation of specific genes upon the transition from one state to another, we calculated for PHO3 cells how much the mean value of the expression level for each considered gene differed upon the observed transitions between clusters (Supplementary Table 2). This yielded for each transition of the Petri net of Figs 6A,7A the x-fold change in the expression level of each gene that occurs upon firing of the transition. These values are used as arc weights in the gene expression modules of the Petri net. There are multiple copies of the gene expression module shown in Fig. 9C which are used to update the expression-level of each considered gene upon firing of each of the Petri net transitions of Figs 6A,7A (see legend to figure Fig. 9C for structural details and functionality of the Petri net module). The expression level of genes in terms of relative mRNA concentration is represented by the number of tokens in the place for its gene-specific mRNA. As each transition from one cluster to another is associated with the differential regulation of quite a number of genes, we use logic transitions and logic places for the gene-specific mRNAs to graphically disentangle the network in order to avoid a confusing crisscross of arcs. The model is constructed based on the x-fold changes in the expression level of each gene that occurs during switching between clusters. An example is provided for transitions that are involved in commitment of PHO3 cells to sporulation (x-fold changes, listed for transitions T28 and T78 in Table 3). By employing gene expression modules in the described form, we obtain not more than a formal model of the phenotype of a cell rather than a mechanistic model representing molecular causality. Nevertheless, we suggest that the phenotype model is useful as it predicts possible single cell trajectories, decomposes complex phenotypes (Table 3) in a formalized manner, and therefore may serve as a scaffold for the computational inference of a model of the molecular causality directly from experimental data [43, 44] (see Discussion).

## 4. Discussion

### 4.1. Petri net representation of single cell trajectories

We have analysed differential gene expression in individual *Physarum polycephalum* plasmodia that were exposed to a differentiation-inducing far-red light stimulus pulse by repeatedly taking samples from the same plasmodial cells. MDS analysis revealed that data points were not evenly spread over the plot but formed clouds, with different patterns obtained for cells with different genotype. Connecting data points from single cell time series showed that cells followed different trajectories in the MDS plot in switching between clouds of data points. These trajectories obviously differed between cells that had or had not been exposed to far-red light and that sporulated or did not sporulate in response to the light stimulus. Although cells responded to a stimulus and eventually were committed to sporulation, their response in terms of differential gene expression was obviously different between individual plasmodial cells, as it was previously reported for the wild type [23]. Again, this behaviour was genotype-dependent. In the non-sporulating mutant PHO57 trajectories of stimulated and not stimulated cells could not even be discriminated.

Hierarchical clustering of the data points obtained for the cells of each genotype, PHO3, PHO57, and the PHO3 + PHO57 heterokaryon, identified significantly different clusters in each of the three data sets. These clusters mapped to corresponding clouds of data points in the MDS plot, confirming that both statistical methods delivered consistent results and assigning discretized states of gene expression to the clouds of data points in the MDS plot. Based on the discretized states, single cell trajectories were evaluated for their time-dependent switching between clusters. From this data set and from averaged differences in gene expression between clusters we reconstructed Petri net models that predict single cell trajectories and spontaneous or light-induced changes in gene expression of sporulating and not sporulating cells.

In order to discuss the biological relevance of our Petri net models, a couple of points should be mentioned. First of all, this type of Petri net is a phenotype model. It abstracts, mimics, and predicts the discretized behaviour of the cells in terms of spontaneous and stimulus-induced gene expression and the associated commitment to sporulation. Clearly, it is not a model that explains the cellular behaviour based on causal molecular mechanisms. However, it makes a systems-oriented, formalized approach to the analysis and decomposition of complex phenotypes, and the model may therefore be useful as a scaffold to reverse engineer the underlying network of molecular mechanisms.

Another critical point that should be noted is that the data sets that were available to construct the Petri net models are still small. With more data analysed the number of clusters and hence the number of places in the Petri net might change and the number of transitions might become larger. Hence the Petri nets shown here just reflect the data sets as they currently are but despite this limitation do provide a proof-of-principle for the Petri net based computational approach.

### 4.2. How does the Petri net model relate to the Waddington quasipotential landscape?

Conrad Hal Waddington introduced the metaphor of an epigenetic landscape to intuitively explain canalization of development and cell fate determination during embryogenesis and its dependence on gene mutations [45]. He thought of cells that make developmental decisions to differentiate into alternative cell types as marbles rolling down a landscape with multiple bifurcating valleys finally ending up in separate valleys that correspond to the alternative states of terminal differentiation. The specific topology of this landscape, according to his metaphor, is determined by an underlying layer of genes with ropes attached that pull the "epigenetic" landscape (the landscape arranged like a sailcloth above the genes) into shape [45]. Alterations to those genes would ultimately reshape the topology with impact on the development of an individual. This is where the term epigenetics originally comes from.

Theoretical considerations of multistability in boolean network models of gene regulatory networks has provided a theoretical basis for a possible molecular interpretation of Waddington’s landscape. It may be understood as a quasipotential landscape (QPL) of a dynamic system of coupled biochemical reactions that form feedback loops. The valleys of the QPL then correspond to different attractors and the altitude of the mountaintops is a measure for the unlikeliness that the system will be in a particular state while the state space of the system is projected into the two-dimensional plane [2, 46-51].

From bacteriophages to mammals, decision-making in the regulation of cell cycle, differentiation, and development is central to a wide variety of developmental phenomena. Indeed, in cases where molecular mechanisms have been elucidated, it could be computationally demonstrated that such phenomena can be readily explained on the basis of bistability and multistability of the underlying biochemical reaction networks [52-62]. Computational approaches have also shown how well-known kinetic mechanisms of interacting biomolecules can result in a quasipotential landscape with multiple attractors and how mechanisms of stem cell differentiation, transdifferentiation, and carcinogenesis can be understood in the context of such a landscape [2, 51, 63-68]. Most recently, a synthetic gene regulatory network of inducible promotors and transcriptional regulators with mutual inhibitions and autoactivations implementated in *Escherichia coli*, was experimentally shown to exhibit quadrastability. Elegant experimental and computational evidence demonstrated that the topology of the resulting quasipotential landscape depends on the connectivity within a network. The connectivity was permutated systematically by adding different combinations of inducers and repressors controling the promotors of the network, switching on and off regulatory interactions between the nodes [69].

The concept of a quasipotential landscape to regulate cell fate determination offers more than just a theoretical framework to explain multistability. It suggests that attractors and the multistability of functional states of the regulatory network of a cell can eventually be reached through several alternative pathways of cellular regulation that are taken and that cells can switch between stable or meta-stable states as a result of stochasticity or transient perturbation of the reaction network [46]. A rugged landscape enabling variable pathways hight have emerged from evolutionary constraints on the versatility, adaptability, and robustness of regulatory networks that have led to the development of highly complex, multistable networks in order to gain a sufficient degree of fitness. Obviously, variable pathways connecting different attractors of a QPL might not only lead to states of disease but also be exploited for rationally-designed medical treatment [67, 70].

Variable pathways for switching between two states of cell differentiation have been directly demonstrated in the unicellular eukaryote *Physarum polycephalum* quite a number of years ago [71]. Morphological intermediates in the form of mitotic figures and cytoskeletal rearrangements that occur during the development of an amoebal cell into a plasmodium were directly observed in the light microscope [71]. At the molecular level, the variability of pathways has also been demonstrated for the development of the *P. polycephalum* plasmodium into fruiting bodies (sporulation) which is another differentiation process that occurs in the course of the life cycle [23].

Can a Petri net as a predictive model for single cell trajectories be helpful to retrive information about the Waddington quasipotential landscape? As biochemical reactions are driven by differences in the chemical potential, statistically significant different single cell trajectories of measured observables (like mRNA concentrations or changes in the abundance or covalent modification of proteins) will ultimately reflect differences in causal mechanisms or processes that influence these observables. The clouds of data points in the MDS plots indicate states of the system that are likely while the white regions indicate states that are less likely and the density of data points, at high sample size, is a measure of the likeliness that the system adopts a certain state. In this work, such relatively likely states have been discretized by hierarchical clustering and modeled as places of a Petri net as shown by the cartoon of Fig. 10. The probabilities that the system transits from one state to the other can be encoded as rate constants assigned to stochastic Petri net transitions. Hence, we assume that single cell trajectories, e.g. as projected into the plane of an MDS plot or as predicted by a corresponding Petri net model, do reflect pathways that are taken through the QPL. Accordingly, we propose that the Petri nets of Figs. 6,7 as executable models do reflect aspects of structure and dynamics of the QPL of a cell of a given genotype. As indicated by the dynamic behaviour and by the T-invariants of the Petri nets derived for strain PHO3 and for the PHO3 + PHO57 heterokaryon, the system can transit between certain states that are meta-stable or even stable. At least some of these states may correspond to basins of attraction. Upon photo-stimulation, stimulus-dependent Petri net transitions are enabled, as the topology of the QPL has changed and the system now moves to a new basin of attraction through states indicated by the Petri net places. Accordingly, responsiveness of the topology of the QPL to differentiation-inducing stimuli in the Petri net framework is modeled in the form of specific transitions that, if enabled, allow the system to proceed to a new state of attraction.

**Figure 10.**
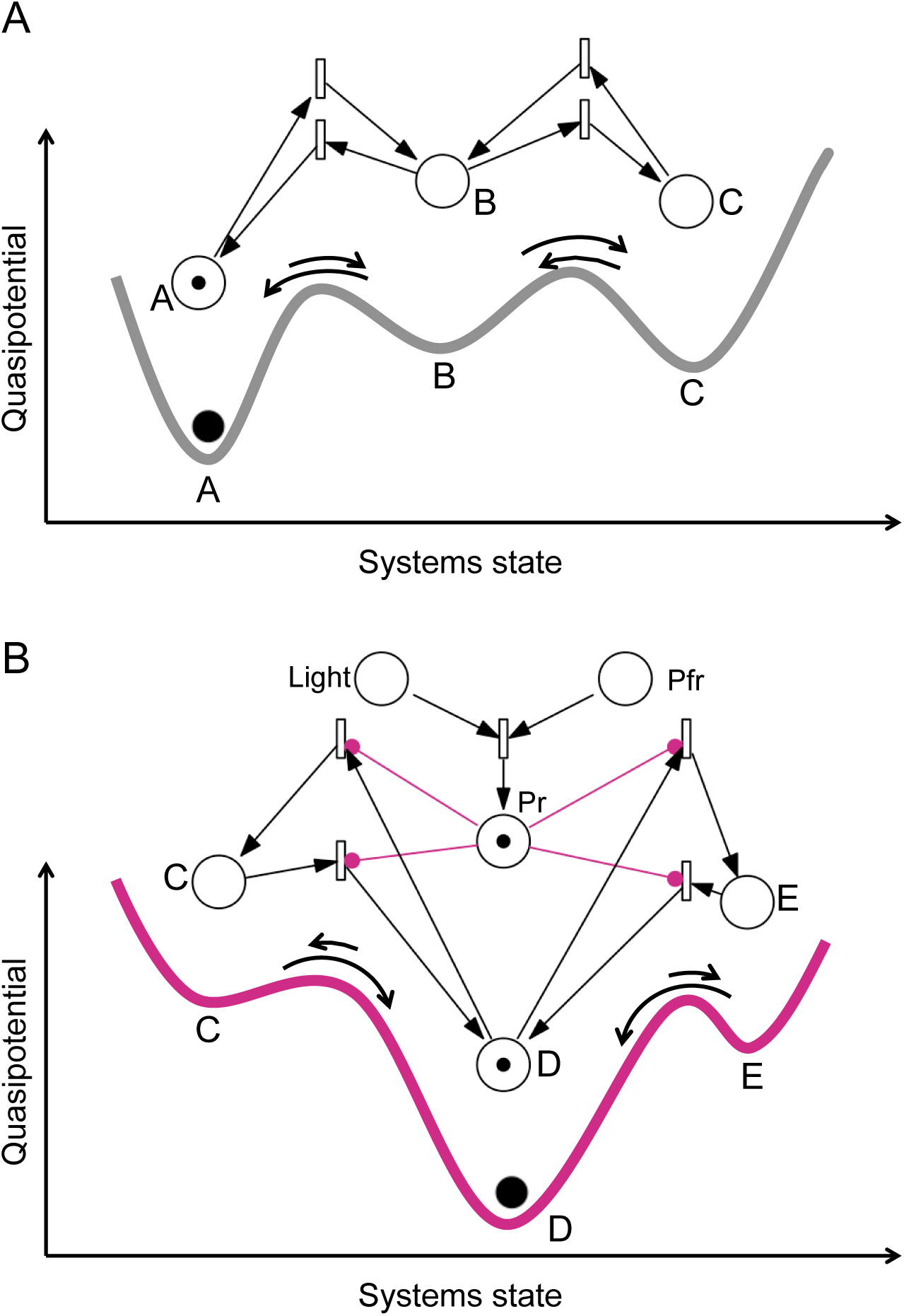
Interpretation of Petri net places and transitions in the context of the Waddington quasipotential landscape. The cartoon suggests that some of the places of the Petri nets of Figs. 6 and 7 may represent states of gene expression (represented by the marble and the token) that correspond or lead to basins of attraction of the Waddington quasipotential landscape. Upon light stimulation, the topology of the quasipotential landscape of unstimulated cells (A) is remodeled (B) so that the system is exposed to new basin(s) of attraction. This might for example occur when photoactivation of the phytochrome photoreceptor alters certain steady states within the regulatory network.

## Acknowledgments

We thank the anonymous reviewers for the careful assessment of our work and for numerous valuable suggestions that lead to a considerable improvement of this manuscript. B.W. was supported by the state of Saxony-Anhalt through the International Max Planck Research School for Advanced Methods in Process and Systems Engineering, Magdeburg.

## Supplementary Information

Supplementary Information provided through the online version of this article:

Supplemental Movie 1

Supplemental Table 1

Supplemental Table 2

